# N439K variant in spike protein may alter the infection efficiency and antigenicity of SARS-CoV-2 based on molecular dynamics simulation

**DOI:** 10.1101/2020.11.21.392407

**Authors:** Wenyang Zhou, Chang Xu, Pingping Wang, Meng Luo, Zhaochun Xu, Rui Cheng, Xiyun Jin, Yu Guo, Guangfu Xue, Liran Juan, Huan Nie, Qinghua Jiang

## Abstract

Severe acute respiratory syndrome coronavirus 2 (SARS-CoV-2), causing an outbreak of coronavirus disease 2019 (COVID-19), has been undergoing various mutations. The analysis of the structural and energetic effects of mutations on protein-protein interactions between the receptor binding domain (RBD) of SARS-CoV-2 and angiotensin converting enzyme 2 (ACE2) or neutralizing monoclonal antibodies will be beneficial for epidemic surveillance, diagnosis, and optimization of neutralizing agents. According to the molecular dynamics simulation, a key mutation N439K in the SARS-CoV-2 RBD region created a new salt bridge which resulted in greater electrostatic complementarity. Furthermore, the N439K-mutated RBD bound hACE2 with a higher affinity than wild-type, which may lead to more infectious. In addition, the N439K-mutated RBD was markedly resistant to the SARS-CoV-2 neutralizing antibody REGN10987, which may lead to the failure of neutralization. These findings would offer guidance on the development of neutralizing antibodies and the prevention of COVID-19.

## 1 INTRODUCTION

Coronavirus disease 2019 (COVID-19), caused by a single-stranded positive-strand RNA virus named severe acute respiratory syndrome-coronavirus 2 (SARS-CoV-2), is a major threat to public health worldwide[1–3]. As of 2 August 2020, over 17 million confirmed cases including 675,060 COVID-19 related deaths were reported worldwide according to the World Health Organization (https://www.who.int/emergencies/diseases/novel-coronavirus-2019/situation-reports). It has been reported that SARS-CoV-2 infects humans through the binding of the homo-trimeric spike (S) glycoprotein to human angiotensin converting enzyme 2 (hACE2), and this infection mechanism for viral entry is also used by SARS-CoV[2, 4]. The surface spike glycoprotein including S1 and S2 subunits is the major antigen of coronaviruses, and S1 binds to host cells whereas S2 mediates viral membrane fusion. The receptor-binding domain (RBD) mediates the binding of the virus to host cells, which is a critical step for viral entry[2, 5–7]. According to the high-resolution crystal structure[8, 9], the receptor-binding motif (RBM) is essential for RBD and contacts highly with hACE2. The structural characterization of the pre-fusion S protein provides atomic level information to guide the design and development of antibody[10].

Neutralizing monoclonal antibodies of the immune system, which play an important role in fighting against viral infections, have been found to target the SARS-CoV-2 RBD and exert neutralization activity by disrupting the virus binding[11–15]. During the virus transmission, alterations of amino acid in the surface spike protein may significantly alter the virus antigenicity and the efficacy of neutralizing antibodies. As SARS-CoV-2 spreads around the world, the mutations in spike protein had been continuously reported[16–20]. There are 930 naturally occurring missense mutations in SARS-CoV-2 spike protein that had been reported in the GISAID database (Supplementary Table S1), and a key mutation from ASP614 to GLY614 (D614G) in SARS-CoV-2 spike protein made the virus more infectious than the original strain[21]. However, most immunogens, testing reagents, and several antibodies for SARS-CoV-2 are based on the S protein of the Wuhan reference sequence (GenBank: MN908947.3). Among coronaviruses, missense mutations had been demonstrated to confer resistance to neutralizing antibodies in MERS-CoV and SARS-CoV[22–24]. In the case of HIV, missense mutations are known to influence envelope glycoprotein expression, virion infectivity[25], alter neutralization sensitivity[26], and confer resistance with neutralizing antibodies[27, 28]. The epidemiological observations have proved the mutations, especially in S protein, provide a plausible mechanism for the increased observed infectivity and bring challenges to antibody development for SARS-CoV-2.

Here, we present findings from molecular dynamics (MD) simulations of the binary complexes of the RBD domain with the common receptor hACE2 and the neutralizing antibody CB6/REGN10987, respectively. The structural basics of N439K-mutated RBD-hACE2 complexes show a new salt bridge and local interaction. The energetic details further indicate that the mutated RBD-hACE2 complexes show higher affinity than the original strain which are mainly attributed to the changes in van der Waals and electrostatic energies of some key residues. On the other hand, N439K reduced the sensitivity to neutralizing antibodies which are mainly attributed to polar solvation and electrostatic interactions. Taken together, the SARS-CoV-2 spike protein with N439K may be more infectious and become resistant to some SARS-CoV-2 neutralizing antibodies.

## 2 RESULTS

### 2.1 The diversity of mutations in SARS-CoV-2 whole-genome sequence

Our trajectory analysis of SARS-CoV-2 mutations in the COVID-19 pandemic was established on 64039 SARS-CoV-2 genome sequences which were downloaded from the Global Initiative for Sharing All Influenza Data (GISAID) database. The whole-genome sequence of SARS-CoV-2 can be divided into two categories according to their function, the open reading frame (ORF) encoding non-structural proteins, and the main structural proteins including spike (S), envelope (E), membrane (M), and nucleocapsid (N). The SARS-CoV-2 sequences were aligned to the Wuhan reference genome with MAFFT[29], then the amino acid changes were identified based on the sequence alignment. In general, the mutation rate of SARS-CoV-2 is a little low[17], and most mutations are significantly concentrated in European and North American populations (Fig. 1A). When observing the frequency of mutations in all genes, the amino acid changes are mostly distributed in ORF1a, ORF1b, N, and S proteins. Interestingly, the ORF3a protein has a superior mutation rate in North American and Oceanian populations accounting for 14.81% and 10.22% of the total protein, respectively (Fig.1A and Supplementary Table S1). The ORF3a protein is also called “accessory protein”, it can convert the environment inside the infected cell and make holes on the infected cell membrane. Hence the mutations on ORF3a protein may give virus the ability to spread efficiently.

**Figure 1.**
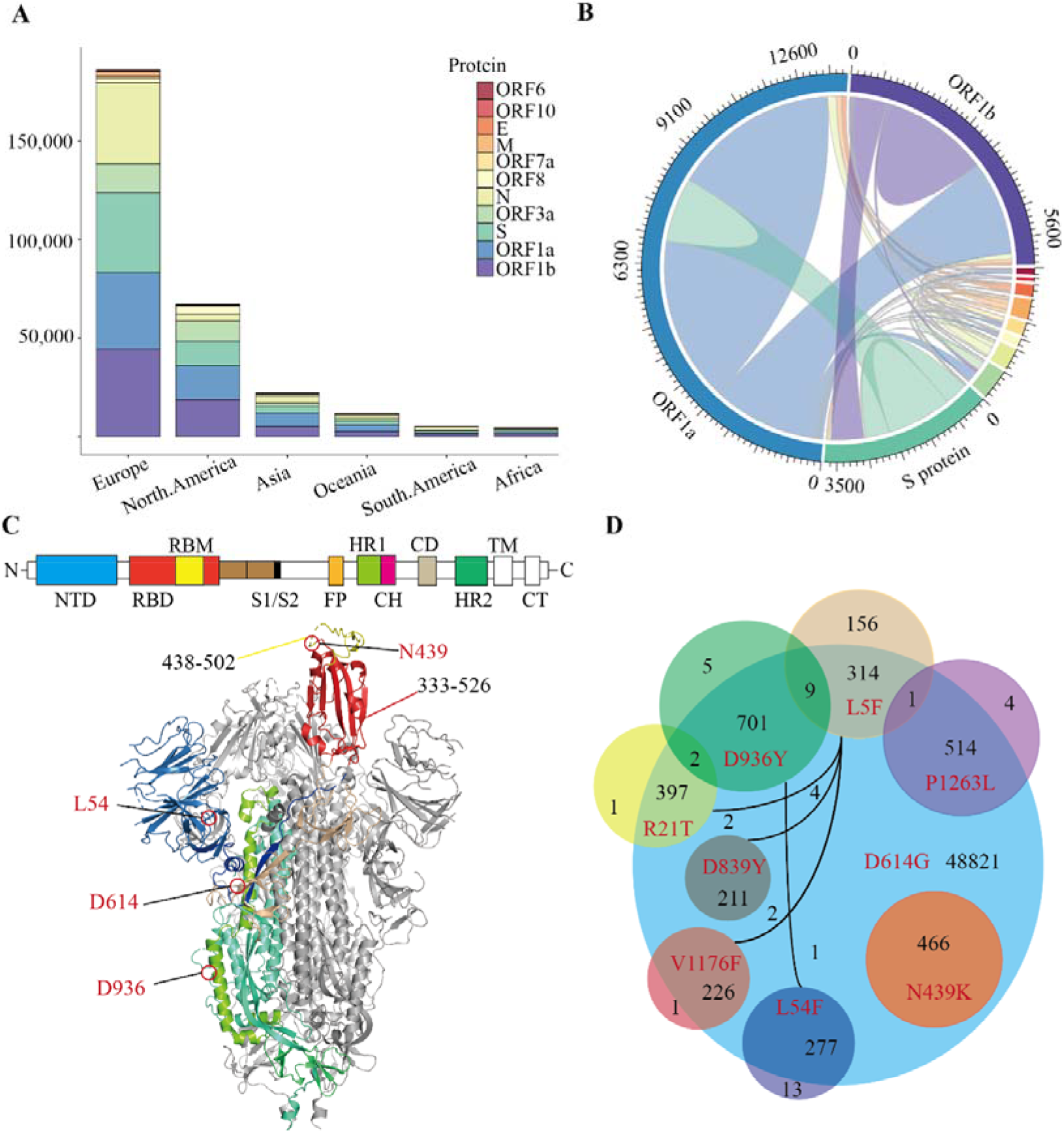
The distribution of Missense Mutations in SARS-CoV-2. (A) A stacked histogram shows all missense mutations of SARS-CoV-2 from different continents of the world. (B) A chord diagram shows the missense mutation shared between different proteins of SARS-CoV-2, and the color scheme is the same as that in Fig.1A. (C) A topology and cartoon representation of SARS-CoV-2 homo-trimeric spike (S) glycoprotein (PDB ID: 6VSB). NTD, N-terminal domain; FP, fusion peptide; HR1, heptad repeat 1; CH, central helix; CD, connector domain; HR2, heptad repeat 1; TM, transmembrane region; CT, cytoplasmic tail; S1/S2, protease cleavage sites; Some important mutation sites and residue intervals were marked on the figure. (D) A Venn diagram of main amino acid mutation sites of the S protein, the number of mutations, and co-mutations were shown.

Co-mutation may affect the cooperation between the various proteins, which may lead to meaningful virus evolution and pose challenges to the development of antibodies. To investigate the co-mutation in SARS-CoV-2, we calculated the co-mutation in all proteins and found most mutants were not single-site mutations (Fig.1B). Since SARS-CoV-2 infects humans through the binding of trimeric spike glycoprotein (PDB ID: 6VSB) to hACE2, then we counted all amino acid variants on S protein (Fig.1C and Supplementary Tab.S1). The variant D614G accounts for 75.92% of 64039 SARS-CoV-2[17], following by D936Y accounts for 1.11%. And more notably, N439K is the most dominant variant in the RBD region, which accounts for 0.72% of all SARS-CoV-2. Furthermore, there are only 16 mutations on S protein with a mutation rate of more than 0.16% (Supplementary Tab.S1), and those mutants occur frequently together with the variant D614G (Fig.1D). All the N439K are included in the D614G interestingly, which is mainly concentrated in Europe since March 2020 (Supplementary Tab.S1). To conclude, the amino acid changes seem to have been accumulated progressively over time, among which N439K is the most dominant variant in the RBD region and should be preferentially employed to characterize the influence of mutations on pathogen evolution.

### 2.2 The N439K-Mutated RBD binds hACE2 with higher affinity than wild type

The amino acid sequence of N439K differs from wild-type in that there is a mutation from Asparagine (N) to Lysine (K) at the 439th amino acid site (Fig.2A). Crystal structures show that the high contact area of SARS-CoV-2 RBD-hACE2 complex is mainly concentrated in the RBM region[30, 31], and denoted as CR1, CR2, and CR3[32]. To investigate whether N439K can influence the interactions between hACE2 and RBD, we exploited 100ns MD simulations of the binary complexes of hACE2 with RBD (both wild-type and N439K). To further verify the convergence of MD simulations equilibrium, we estimated the root mean square deviations (RMSD) of backbone atoms relative to the corresponding crystal structure, and the wild complex had a relatively smaller average RMSD than the N439K-mutated complex (2.2 vs. 2.4 Å) (Fig.2B and Tab.1). The structural compactness of each system was elucidated by estimating the radius of gyration (R_g_), the average R_g_ values were similar for all the same systems (Tab.1). We also calculated the solvent accessible surface area (SASA) which indicates the solvent exposure degree, and SASA values of wild and mutant complexes were 35716 Å^2^ and 36121 Å^2^, respectively (Tab.1 and Supplementary Tab.S2). To further investigate the detailed residual atomic fluctuations of structural elements in wild and mutant complexes, we computed the root mean square fluctuations (RMSF) for protein C_α_ atoms and provided insights into the structural fluctuation and flexibility. Although the flexibility patterns of residues in the N439K-mutated RBD-hACE2 complex display similar fluctuations with wild-type complex (Fig.2C), certain regions of the two complexes show differences in flexibility. The wild-type complex showed slightly high fluctuations on hACE2, while the N439K-hACE2 complex shows much more fluctuation in the RBD domain. Increment of RMSF values of the N439K RBD domain confirmed that the flexibility changes were significantly strengthened by the 439th amino acid change. This fluctuation of the variant N439K might induce structural rearrangements of the SARS-CoV-2 RBD-hACE2 complex, possibly leading to a stronger binding ability.

**Figure 2.**
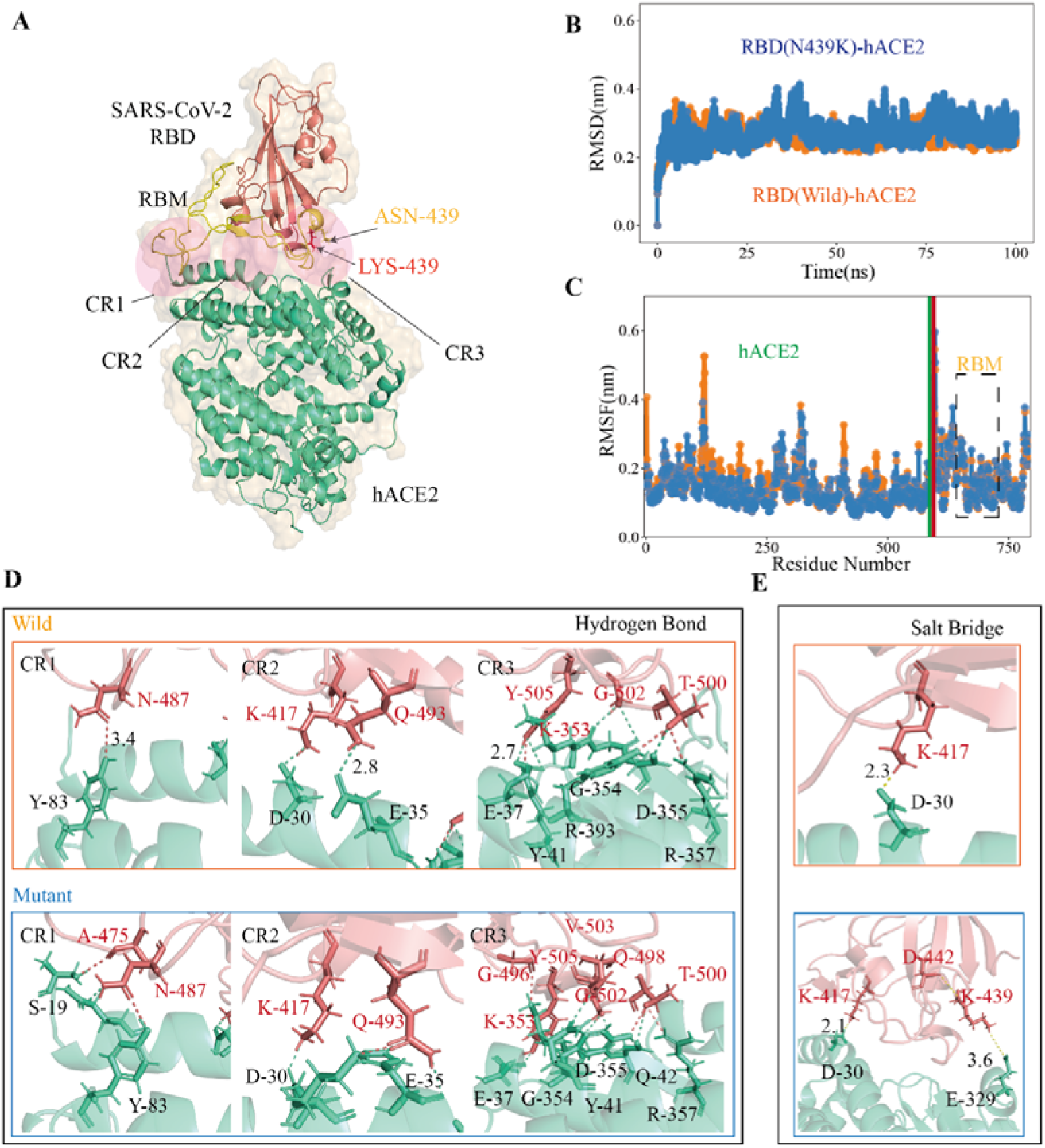
The RBD-hACE2 Interaction Profile of MD Simulations. (A) The structure of SARS-CoV-2 RBD-hACE2 (PDB ID: 6M0J). SARS-CoV-2 RBD core is shown in deepsalmon and its interacting hACE2 is colored greencyan, and RBM of RBD is represented in yellow. Receptor-ligand interface includes the N-terminal (CR1), the central region (CR2) and the binding loop (CR3), ASN439 (yellow) to LYS439 (red) in CR3. (B) The RMSDs of the backbone atoms of both RBD-hACE2 complexes, the RBD-hACE2 is colored orange and RBD(N439K)-hACE2 is colored blue. (C) The RMSFs of *C_α_* atoms of both RBD-hACE2 complexes, where RBD-hACE2 is colored orange and RBD(N439K)-hACE2 is colored blue. (D&E) Hydrogen bonds and salt bridges are shown as dotted lines, RBD (deepsalmon), and hACE2 (greencyan) residues are described as sticks, respectively. The orange box indicates wild-type RBD-hACE2 and mutant-type RBD-hACE2 is a blue box.

**Table 1.**
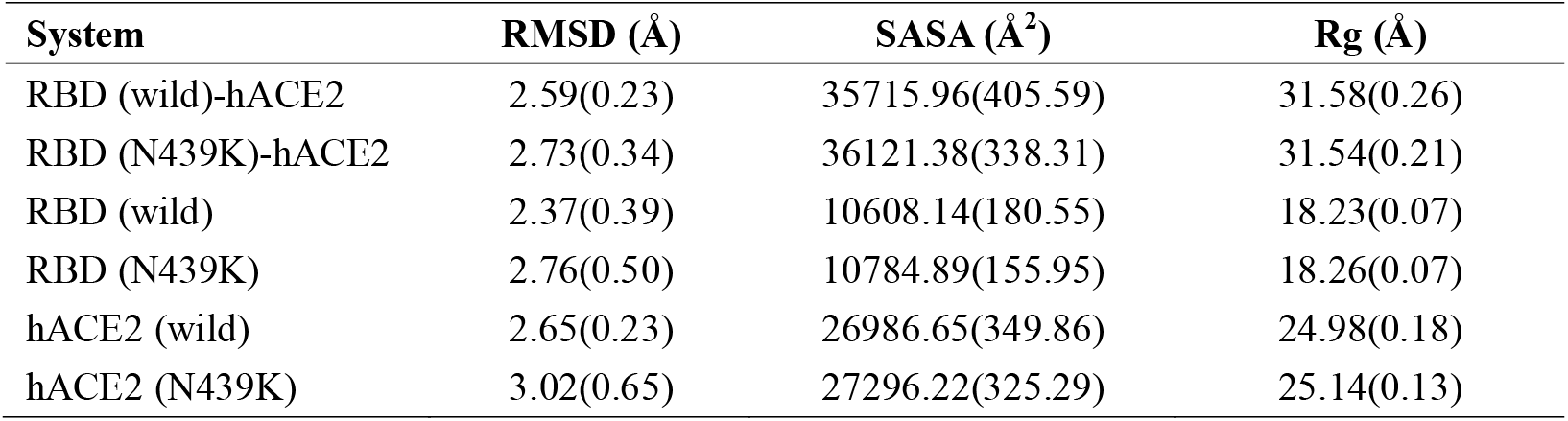
The average RMSD, solvent accessible surface area (SASA), and radius of gyration for the simulated systems. Standard errors of the mean (SEM) are provided in parentheses.

| System | RMSD (Å) | SASA (Å<sup>2</sup>) | Rg (Å) |
| --- | --- | --- | --- |
| RBD (wild)-hACE2 | 2.59(0.23) | 35715.96(405.59) | 31.58(0.26) |
| RBD (N439K)-hACE2 | 2.73(0.34) | 36121.38(338.31) | 31.54(0.21) |
| RBD (wild) | 2.37(0.39) | 10608.14(180.55) | 18.23(0.07) |
| RBD (N439K) | 2.76(0.50) | 10784.89(155.95) | 18.26(0.07) |
| hACE2 (wild) | 2.65(0.23) | 26986.65(349.86) | 24.98(0.18) |
| hACE2 (N439K) | 3.02(0.65) | 27296.22(325.29) | 25.14(0.13) |

Subsequently, we performed the hydrogen bond (H-bond) and salt bridge analyses based on the trajectories in 100ns MD simulations. The results exhibited distinct epitope features between wild-type and mutant, although they can both engage hACE2. It was observed that the N439K-mutated RBD-hACE2 complex can form more hydrogen bonds than the wild-type (Fig.2C). Hydrogen bonds from the N-terminal (CR1) to the central region (CR2) of the interface were almost the same during MD simulations. However, three additional main-chain hydrogen bonds forms at Gly496, Gln498, and Val503 in the binding loop (CR3), causing the ridge to take a more compact conformation and the loop to move closer to hACE2 (Fig.2C). At the SARS-CoV-2 RBD-hACE2 interface, a strong salt bridge[33] between Lys417 of the RBD and Asp30 of hACE2 had been confirmed[8] (Fig.2D). The mutation of N439K in the RBD formed a new salt bridge with Glu329 of hACE2 (3.6 Å), where Arg426 of SARS-CoV RBD also formed a salt bridge at the same position[9], and a weak salt bridge between Lys439 and Asp442 of SARS-CoV-2 RBD (Fig.2D). Burial of these salt bridges in hydrophobic environments on virus binding would enhance their energy owing to a reduction in the dielectric constant.

To estimate the stabilization of binding systems, We also calculated the molecular mechanics Poisson-Boltzmann surface area (MM-PBSA) in both wild and mutant SARS-CoV-2 RBD-hACE2 complexes.[34]. The combination of MD simulation with MM-PBSA incorporated conformational fluctuations and entropic contributions to the binding energy. The components in the binding free energy of RBD-hACE2 complexes include ΔE_vdw_, ΔE_elec_, ΔG_pol_ and ΔG_apol_. Overall, the binding free energy (ΔG_bind_) decomposition analysis divulged into various free energies (Fig.3B and Supplementary Tab.S3). The binding energy ΔG_bind_ (−1526.17+/−133.13kj/mol) of the N439K-hACE2 was higher in magnitude as compared to the wild-type RBD-hACE2 (−1084.06+/−80.23kj/mol) (Fig.3A), in which the electrostatic energy between wild and mutant types had a significant difference. Meanwhile, we calculated binding energy between RBM (residue 438 to 502) and hACE2 using MM-PBSA, and binding energies of the two complexes were −860.94+/−99.45kj/mol and −398.06+/−111.76kj/mol, respectively (Fig.3C and Supplementary Tab.S2). Comparing the binding free energy of RBD-hACE2 and RBM-hACE2, it should be noted that the changes in energy are mainly concentrated in the RBM-hACE2 region.

**Figure 3.**
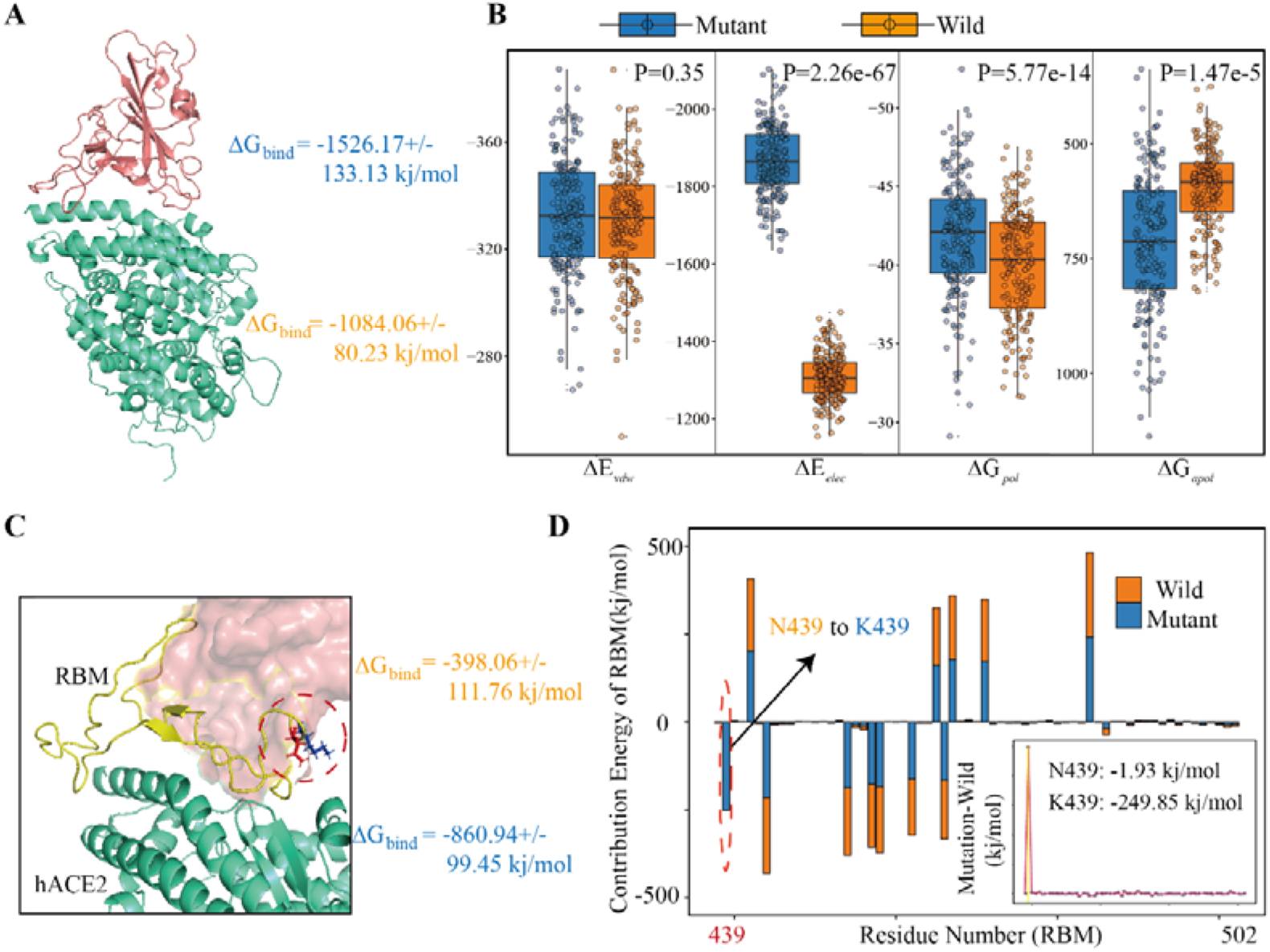
Energy Components for Binding Free Energy of Both Wild and Mutated RBD/RBM-hACE2 complexes. (A) The binding free energies for SARS-CoV-2 RBD-hACE2 (including wild type and variant N439K) using MM-PBSA. Total binding energy (ΔG_bind_). (B) Energy components for the binding free energy of RBD-hACE2. The intermolecular van der Waals (ΔE_vdw_); Electrostatic interactions (ΔE_elec_); Polar solvation energy (ΔG_pol_) and apolar (non-polar) solvation energy (ΔG_apol_). (C) The binding free energies for SARS-CoV-2 RBM-hACE2 (including wild type and variant N439K), using orange for wild-type and blue for mutant type. (D) Decomposition of ΔGbind into contributions from individual residues (438-502) of RBM before and after mutation. All units are reported in kj/mol.

Subsequently, we explored the critical residues involved in the RBM-hACE2 binding by performing the per-residue decomposition of binding free energy (Fig.3D), and the contribution energy of per-residue to the total binding energy was compared before and after mutation. It is evident from the line graph (Fig.3D) that Lys439 hotspot residue show more contribution to binding free energy than ASN439 and its contribution energy changed from −1.93kj/mol to −249.85kj/mol (Fig.3D). Taken together, comparing the interaction interfaces of the N439K-mutated and wild RBD-hACE2 complexes reveals the change from ASN439 to LYS439 might result in a tighter association because of the new salt bridge formation and higher affinity.

### 2.3 N439K became resistant to SARS-CoV-2 neutralizing antibody REGN10987

To investigate the antigenicity of the N439K mutant, we exploited 100ns MD simulations of the binary complexes of hACE2 with neutralizing monoclonal antibodies REGN10987 and CB6 complexes. Specific neutralizing mAbs can prevent the virus from binding to hACE2 by neutralizing SARS-CoV-2. Although most of these highly binding sites are in RBM, the two neutralizing antibodies show different high contact regions with RBD (Fig4.A), REGN10987 binds mainly to CR2 and CR3 regions of RBD where the N439K located, and the main binding regions of CB6 are CR1 and CR2.

**Figure 4.**
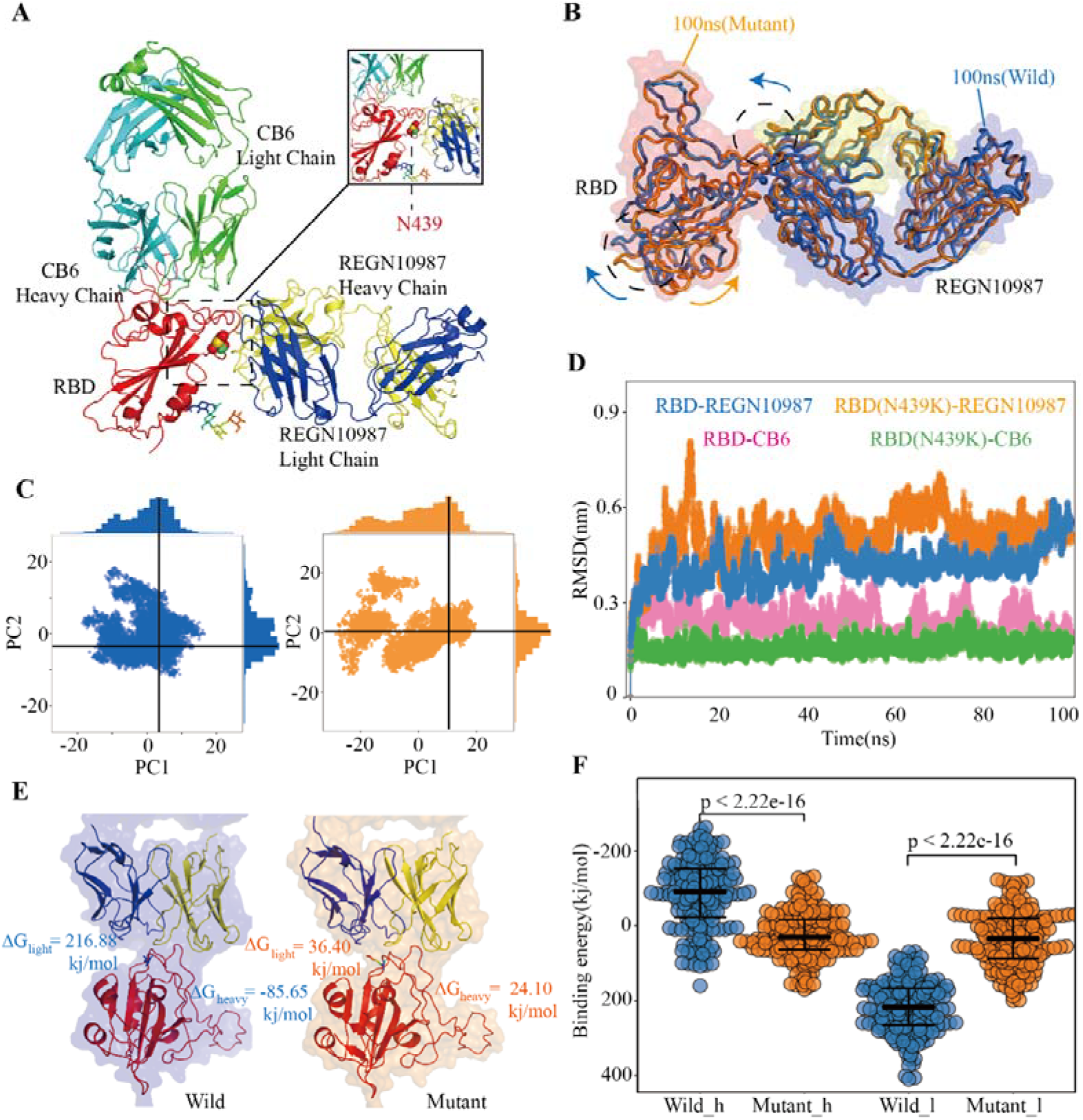
Structural and Energetic Details of Both Wild and Mutant RBD-mAbs Interactions. (A) Crystal structures of RDB-CB6/REGN10987 complexes, the RBD is colored red, CB6 heavy and light chains are represented as marine and green respectively, REGN10987 heavy chain is colored yellow and light is blue, and the 439 residues are described as the sphere. (B) Characteristic dynamic fluctuations of both RBD-REGN10987 and RBD(N439K)-REGN10987 complexes. Mutant-type (100ns) and wild-type (100ns) are colored by orange and blue, respectively. (C) Dynamic conformations are projected on to the principal vectors (PC1 and PC2). Red and blue indicate mutant-type and wild-type 100ns MD trajectories respectively. (D) The RMSD of the receptor-binding motif in four complexes during the 100-ns MD simulations. (E) The binding free energies for both complexes of the mAb REGN10987 (including heavy and light chains), the color schemes are the same in Fig.4A and Fig.4C. (F) The binding free energies of 200 configurations at an interval of 100ps from the last 20ns simulations. The t-test was conducted to check the statistical significance of the difference between two systems of binding free energies. A p-value of <0.05 indicates that the difference is statistically significant (95% confidence interval). The color scheme is the same as that in Fig.4C

After the 100ns MD simulation, the final conformational distribution indicated the configuration fluctuations of RBD were mainly concentrated in the CR3, whereas the REGN10987 antibody was relatively stable (Fig.4B). Overall, SARS-CoV-2 RBD and REGN10987 undergo symmetric twist and antisymmetric hinge-bending motions about the axis of the N-terminal helix, corresponding to the lowest quasi-harmonic models, PC1, and PC2, from principal component analysis (PCA)[35]. Although the overall conformational dynamics were almost the same between the wild and mutant complexes, the average conformation of the variant N439K-REGN10987 structure has changed relative to wild complex when the dynamic configurations are projected onto the two principal vectors (Fig.4C). Interestingly, the comparison of the crystal structures of two complexes shows a small but significantly conformational difference, perhaps due to crystal packing restraint. From results of RMSD, the wild and mutant systems had reached a stable state from their respective MD trajectories, CB6 systems had lower average RMSD values than the REGN10987 complexes, and a higher RMSD was calculated in N439K-REGN10987 than wild-type (5.1 vs. 4.1 Å) (Fig.4D). The MD simulation results of the CB6-RBD complex can be found in Supplementary materials.

To understand the biophysical basis of the molecular recognition of RBD-mAbs complexes, the MM-PBSA scheme was employed. To further explain the ability of heavy and light chains of mAbs to neutralize viruses, we had taken 200 structures from the stable region of the last 20ns trajectories to calculate binding free energy with RBD respectively. It revealed that the estimated binding free energy of wild-type CB6-RBD complexes (−129.73kj/mol) is higher than N439K-mutated CB6-RBD complexes (−51.28kj/mol) in the heavy chain, the unfavorable contribution from ΔG_pol_ (491.56kj/mol) was relatively higher compared to mutant type (703.82kj/mol) (Tab.2 and Supplementary Tab.S4). Overall, our research suggested that the N439K reduced the sensitivity to the CB6 mAb. Subsequently, we have investigated the binding free energy of REGN10987-RBD complexes, the ΔG_bind_ of N439K-mutated REGN10987-RBD complexes (24.10kj/mol) was found to be lower (P=2.22e-16) than wild-type (−85.65kj/mol) in the heavy chain (Fig.4F). Similarly, the light chain had a little difference in binding free energy with RBD, but both systems could not form an effective combination (Fig.4E and Tab.2). Furthermore, the present 100ns trajectory revealed that polar solvation and electrostatic interaction might be the main factors to lose the ability of the REGN10987 antibody to neutralize COVID-19 (Tab.2).

**Table 2.**
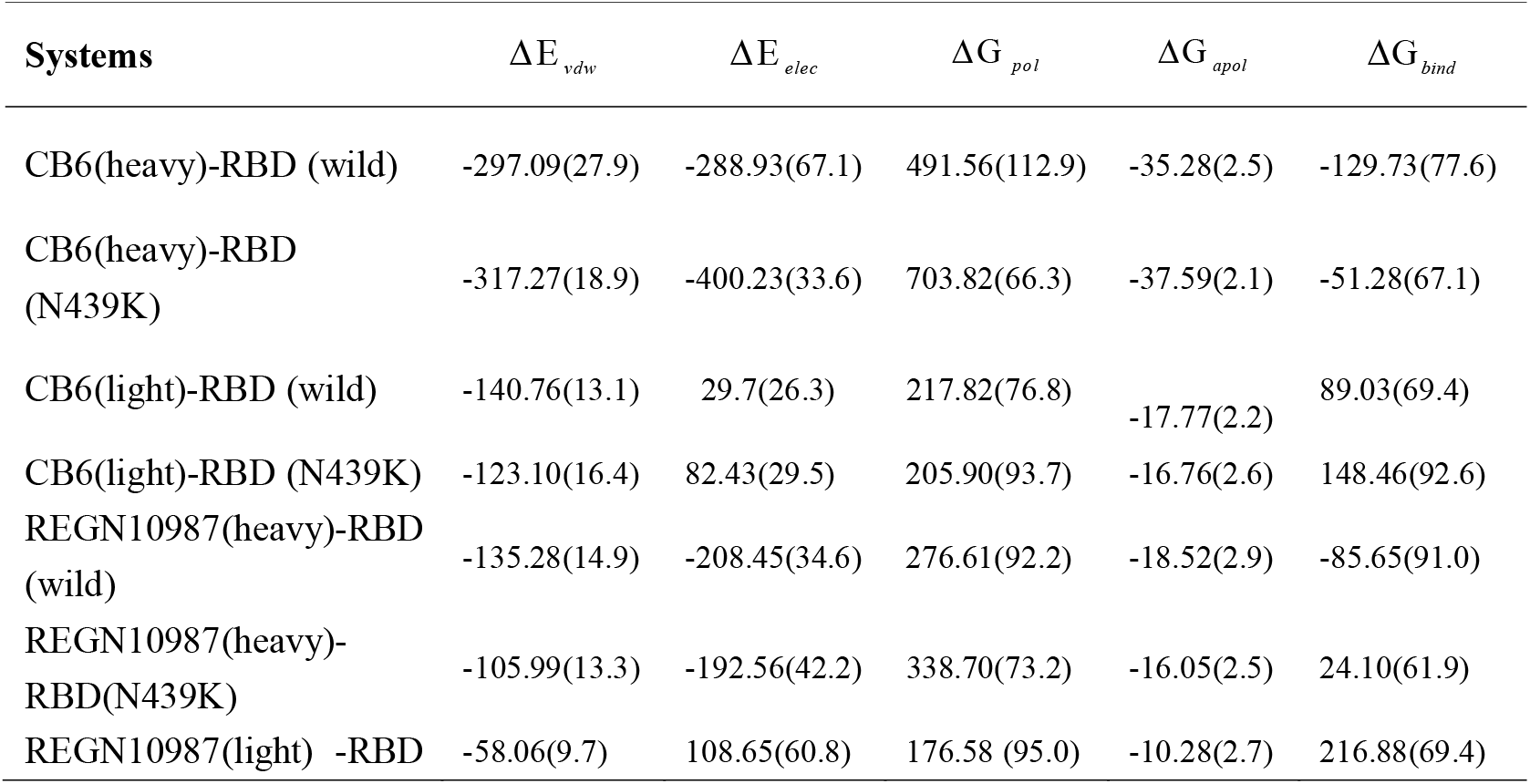
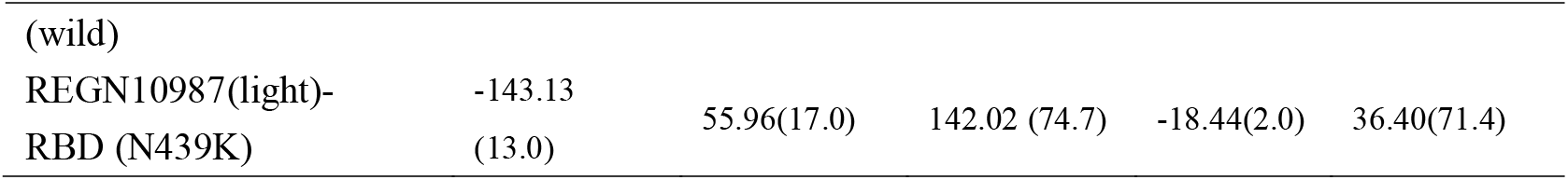
Energetic components of binding energy for SARS-CoV-2 RBD-mAbs complexes (kj/mol). Standard errors of the mean (SEM) are provided in parentheses.

## 3 DISCUSSION

The recent outbreak of COVID-19 has caused a severe strain in the public health system in many countries. The SARS-CoV-2 virus is expected to continue evolving in human populations. Close monitoring of circulating virus strains is of unquestionable importance to inform research and development of antibodies and therapeutics. Herein, we have studied the mechanism of binding of RBD-hACE2 and RBD-mAbs by using an atomistic molecular dynamics simulation of 100ns in conjunction with molecular mechanics/Poisson-Boltzmann surface area (MM-PBSA) scheme. Our study shows that the variant N439K influences the affinity of both RBD-hACE2 and RBD-mAbs complexes which is favored by the intermolecular van der Waals, electrostatic interactions and polar solvation free energy. Our finding may facilitate the development of decoy ligands and neutralizing antibodies for suppression of viral infection.

It had been reported most amino acid variants in RBD, including the N439K, were less infectious[21]. However, MD simulations and MM-PBSA results showed that the binding ability of the N439K-mutated RBD with hACE2 was enhanced, in other words, viral infectivity was enhanced significantly. Subsequently, we will exploit the contradictory findings from the following aspects. Firstly, in terms of method, MD simulations and MM-PBSA were widely used in drug screening, structural analyses, and mutagenesis studies. For instance, a delicate balance of specific and non-specific hydrogen-bonding and hydrophobic networks helped elucidate the similarities and differences in receptor binding between SARS-CoV-2 and SARS-CoV. Meanwhile, these methods can facilitate the ease of finding antiviral drugs for SARS-CoV-2[36]. In the case of HIV-1 protease, these were applied to find the loss of flexibility in the mutants which can inhibit protease activity[37].

Subsequently, the replacement of ARG426 (SARS-CoV RBD) with ASN439 (SARS-CoV-2 RBD) appeared to weaken the interaction by eliminating one important salt bridge with ASP329 on hACE2 and reduced the affinity[8, 9]. Besides, the variant N439K only one amino acid change from ASN439 to LYS439 (basic amino acid) may result in a tighter association with hACE2 at the RBD-hACE2 interface because of the formation of a new salt bridge. What’s more, we also found that the electrostatic interaction may be the main reason for increasing the binding capacity, which was consistent with the formation of the salt bridge. Finally, we interestingly found the variant N439K was completely included in the D614G samples (Fig.1D and Supplementary Tab.S1). D614G was significantly more infectious which had been confirmed, yet there is no report about the influence of this co-mutation on viral infectivity[17]. To further validate the reliability of MD simulations, we also exploited another two cryo-EM structures (PDB ID 6LZG and 6VW1) to simulate and found that the variant N439K still enhanced the infectivity of COVID-19.

Almost all natural variants that affected the reactivity to neutralizing mAbs were located in the RBD region, we only selected the N439K variant in the RBD region with the most frequent mutations[11, 13, 14]. The CB6 mAb still could neutralize N439K mutated COVID-19, but N439K was markedly resistant to the REGN10987 mAb.

The SARS-CoV-2 virus is expected to continue evolving, some mutations only appear in a certain period and other mutations may appear in an unpredictable way. It is necessary to continuously analyze the SARS-CoV-2 virus according to the mutation frequency and time pattern. Meanwhile, COVID-19 antibodies are developed based on the Wuhan reference genome, it is necessary to consider the impact of different mutations on the effectiveness of neutralizing antibodies. In summary, we have counted the amino acid changes in 64039 COVID-19 and laid special stress on analyzing the infectivity of the variant N439K in the SARS-CoV-2 virus and the effectiveness of well-characterized neutralizing mAbs using MD simulations. The variant N439K enhanced the infectivity of the virus with hACE2 and altered antigenicity to some neutralizing mAbs. Taken together, our findings shed light on the influence of N439K Variant on SARS-CoV-2 infection efficiency and antigenicity.

## 4 Materials and Methods

### 4.1 Multi-sequence Alignment and Structure Preparation

In this study, SARS-CoV-2 sequences were aligned against the Wuhan reference genome (GenBank: MN908947.3) using MAFFT[29]. The wild-type complexes were directly downloaded from the Protein Data Bank (PDB ID: 6M0J, 6XDG, 7C01) and removed the water molecules using VMD[38] software. For mutated complexes, firstly wild-type hACE2 and mAbs structures were obtained by removing the RBD structures from RBD-ACE2 and RBD-mAbs complexes respectively. Then the mutated SARS-CoV-2 RBD (N439K) structures were constructed using homology modeling SWISS-MODEL[39]. Finally, the mutated SARS-CoV-2 RBD was aligned with hACE2 and mAbs based on the RBD-hACE2 and RBD-mAbs structures using PyMOL software. All structural figures were generated utilizing PyMOL software (https://pymol.org/2/).

### 4.2 MD Simulation

For each system, MD simulations were performed using GROMACS 5.14.0, the latest CHARMM36 force field was selected, an explicit solvent model was used by an explicit TIP3P (three-point charge) model[40]. Then complexes were solvated in a rectangular periodic box and had a 10 Å buffer distance from along each side, then a suitable number of Na^+^ and CL^−^ ions were added to neutralize the whole system and mimicked a salt solution concentration of 0.15M. Specifically, the following three stages of MD simulations were performed before the production simulation: (1) A full 50,000-step energy minimization was performed with all atoms unrestrained. (2) Each complex was equilibrated with NVT (No. of particles, Volume, and Temperature) ensemble which temperature of the per-solvated system increased from the 0 to 310K gradually for 1ns. (3) Subsequently, a 50,000,000-step (100ns) production run was performed after 1ns NPT using GROMACS. During the MD simulations, the cutoff distances of 12 Å for the long-range electrostatic through the Particle Mesh Ewald (PME)[41]method and van der Waals interactions were used, and the time step was set to 2 fs in all MD simulations. The SHAKE algorithm was employed to calculate the bonds involving hydrogen atoms. Further analyses[42] (RMSD, RMSF, distances, angles, hydrogen bond calculations, and salt bridge calculations) were performed based on the resulting trajectories by GROMACS tools. For the H-bond interaction analysis, the two limiting factors are adopted as follows: (1) donor-acceptor distance is ≤ 3.5 Å, and (2) donor-hydrogen-acceptor angle is ≥120°.

### 4.3 MM-PBSA Calculation

Binding free energies of RBDs with hACE2 and mAbs were calculated using MM-PBSA program, which is the sum of enthalpy (H) and entropy (S) parts. For each binding complex, 200 configurations were taken at an interval of 100ps from the last 20ns simulations. MM-PBSA was used to calculate the polar solvation energies, nonpolar solvation energies and calculates the free energy of the complex (the binding free energy of the receptor with ligand in a solvent medium). The general expression of the term is

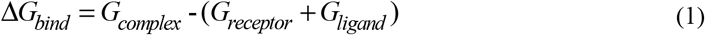

Where G_complex_ is the total free energy complex, and G_receptor_ and G_ligand_ are total free energies of the isolated receptor and ligand in solvent, respectively.

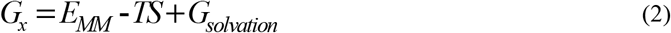

Where x is the receptor or ligand or complex. E_MM_ is the average molecular mechanics’ potential energy in a vacuum. TS refers to the entropic contribution to the free energy in a vacuum where T and S denote the temperature and entropy. G_solvation_ is the free energy of solvation.

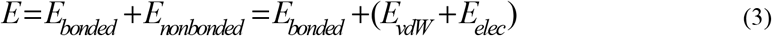

Where E_bonded_ is bonded interactions consisting of bond, angle, dihedral, and improper interactions. E_nonbonded_ includes both electrostatic and van der Waals interactions which are depicted using a Coulomb and Lennard-Jones potential function, respectively. In addition, the free energy of solvation (the energy required to transfer a solute from a vacuum into the solvent) has been calculated including polar and nonpolar solvation energies, it that can be depicted as

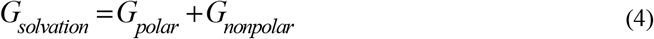

Where G_polar_ and G_nonpolar_ are the electrostatic and non-electrostatic contributions to the solvation free energy, respectively. To check the statistical significance between the two complexes, P-value was calculated using t-test.

## 7 Acknowledgements

This work was funded by the National Natural Science Foundation of China (Nos. 61822108, 62041102 and 62032007 to Q.J.), and the Emergency Research Project for COVID-19 of Harbin Institute of Technology (No. 2020-001 to Q.J.)

## 8 Author contributions

Q.J. and H.N. conceived the project. W.Z. and C.X. collected genome sequences and performed multiple sequence alignment. C.X, Q.J., W.Z, P.W., Z.X., R.C., X.J., G.X., Y.G.,G.X and L.J contributed to molecular dynamics and binding free energy simulations. Q.J., H.N., W.Z. and C.X. wrote the manuscript.

## 10 SUPPLEMENTARY MATERIAL

**Supplementary Tables**

Table S1. Statistics of the mutations of SARS-CoV-2 from GISAID database.

Table S2. Details on both RBD-hACE2 complexes for all simulated systems.

Table S3. Energetic components of binding free energy for RBD/RBM-hACE2 complexes.

Table S4. Binding free energy components for RBD-mAbs complexes.

